# MSICKB: A Curated Knowledgebase for Exploring Molecular Heterogeneity and Biomarker Discovery in Microsatellite Instability Cancers

**DOI:** 10.1101/2025.11.22.689859

**Authors:** Bairong Shen, Yuxin Zhang

## Abstract

**Background:** Microsatellite instability (MSI) is a critical form of genomic instability and a key biomarker for predicting patient prognosis and response to immunotherapy in cancer. However, significant heterogeneity in clinical outcomes exists even among patients with MSI cancers of the same type, suggesting the presence of distinct molecular subtypes and the need for more personalized biomarkers. The systematic exploration of this heterogeneity is currently hindered by the lack of a specialized, multi-dimensional data resource. This raises a critical scientific question: building upon the established link between MSI status and therapeutic efficacy, how can we leverage a multi-dimensional, deeply curated data resource to precisely characterize the molecular heterogeneity of MSI cancers and discover personalized biomarkers for precision oncology?

**Methods:** To address this question, we developed the Microsatellite Instability Cancer Knowledgebase (MSICKB). We systematically curated MSI-related data from 492 publications spanning from 1997 to 2024, covering 31 cancer types. The curated data encompasses four key dimensions: Genetic & Molecular Features, Clinicopathological Features, Therapeutic Response, and Prognostic Factors. Furthermore, we constructed and analyzed a comprehensive gene-disease network to systematically characterize the molecular landscape of MSI cancers.

**Results:** MSICKB(http://www.sysbio.org.cn/MSICKB/) curates a total of 1,382 MSI-related features from 492 publications, covering 31 cancer types. The knowledgebase includes four major categories: Genetic & Molecular Features, Clinicopathological Features, Prognostic Factors, and Therapeutic Response. An application analysis of the curated data revealed that the constructed gene-disease network exhibits a scale-free-like topology, indicating a hierarchical organization governed by a few highly connected hub nodes. Further investigation of the 14 identified hub genes showed significant functional enrichment in core MSI-related pathways, including DNA mismatch repair, immune checkpoint regulation, and key oncogenic signaling pathways.

**Conclusion:** MSICKB provides an integrated, multi-dimensional data resource that facilitates the exploration of the molecular heterogeneity landscape and common regulatory principles within MSI cancers. It serves as a valuable platform for identifying novel molecular subtypes and biomarkers. In the future, the continuous evolution of MSICKB will support the development of clinical prediction models, assist in the advancement of intelligent medicine, and ultimately promote the progress of precision diagnostics and therapeutics for MSI cancers.

## Introduction

Genomic instability is a fundamental hallmark of cancer, contributing to the accumulation of mutations that drive tumorigenesis(1). A key manifestation of this instability is **microsatellite instability (MSI)**, a hypermutable phenotype resulting from a defective DNA Mismatch Repair (dMMR) system(2). Originally identified in hereditary non-polyposis colorectal cancer (HNPCC), or Lynch syndrome(3), MSI is now recognized across a wide spectrum of malignancies, including gastric, endometrial, and pancreatic cancers(4,5).

The clinical significance of MSI status has become increasingly profound. It serves as a crucial **prognostic biomarker**, with MSI-High (MSI-H) colorectal cancer patients often exhibiting a more favorable prognosis compared to their microsatellite stable (MSS) counterparts(6). More importantly, MSI status has emerged as a premier **predictive biomarker** for the efficacy of immune checkpoint inhibitors (ICIs). The high mutational load in MSI-H tumors generates a rich repertoire of neoantigens, rendering them highly immunogenic and particularly susceptible to therapies targeting the PD-1/PD-L1 axis(7). This has led to the landmark tissue-agnostic approval of pembrolizumab for any solid tumor with an MSI-H/dMMR status, heralding a new era of precision oncology(8).

Despite the groundbreaking success of ICIs in MSI-H cancers, a significant clinical challenge persists: **the considerable heterogeneity in patient response**. A substantial portion of patients with MSI-H tumors do not respond to ICI therapy, while others may experience rapid progression or develop acquired resistance(9). This variability strongly suggests that MSI is not a monolithic entity, but rather a complex phenotype influenced by a diverse array of underlying genetic alterations, tumor-specific contexts, and interactions within the tumor microenvironment.

However, a systematic exploration of this molecular heterogeneity is currently hampered by a major bottleneck: **the lack of a centralized, multi-dimensional data resource dedicated to MSI cancers**. While large-scale genomic databases like TCGA and ICGC provide invaluable raw data, they often lack curated, feature-level annotations for clinical outcomes, therapeutic responses, and specific clinicopathological details across a wide range of studies. Consequently, researchers face the arduous task of manually sifting through hundreds of publications to piece together a comprehensive view, severely limiting the potential for large-scale integrative analysis and the discovery of more refined, personalized biomarkers.

To address this critical gap, we have developed the **Microsatellite Instability Cancer Knowledgebase (MSICKB)**, a manually curated and comprehensive repository designed to facilitate the systematic study of MSI cancer heterogeneity. By systematically integrating data from 492 publications, MSICKB consolidates information across four key dimensions: Genetic & Molecular Features, Clinicopathological Features, Therapeutic Response, and Prognostic Factors, covering 31 cancer types.

Furthermore, to demonstrate the utility of this integrated resource for generating novel biological insights, we conducted an application case study. We constructed and analyzed a comprehensive gene-disease network based on the data curated in MSICKB. This analysis allowed us to characterize the topological properties of the MSI-associated molecular network and identify a panel of hub genes in cancers. This work not only provides the scientific community with a valuable, centralized resource but also presents a novel, network-based approach to uncovering the key players driving the complex biology of MSI cancers.

## Methods

### Literature Curation and Data Acquisition

#### Systematic Search Strategy and Study Selection

To construct a comprehensive knowledgebase of features related to microsatellite instability (MSI) in cancer, we performed a systematic literature search of the PubMed database. Our search strategy was designed for maximum sensitivity, employing a free-text search within the title and abstract fields to capture relevant studies that might not be consistently indexed with MeSH terms. The query, executed up to May 12, 2024, was structured as: (MSI[ti] or microsatellite instability[ti]) AND (Cancer[tiab] OR tumour[tiab] OR tumor[tiab] OR neoplasm[tiab] OR carcinoma[tiab] OR lymphoma [tiab] OR Retinoblastoma[tiab] OR melanoma[tiab] OR Seminoma[tiab] OR Nephroblastoma[tiab] OR Osteosarcoma[tiab] OR Sarcoma[tiab] OR Neuroblastoma[tiab] OR mesothelioma[tiab] OR Cholangiocarcinoma[tiab] OR Adenocarcinoma[tiab] OR keratoacanthoma[tiab]).

This initial search yielded 3,314 potentially relevant publications. We then applied a multi-stage screening process (Figure 1). First, we excluded 356 non-primary research articles (e.g., reviews, editorials, case reports) and non-English publications, leaving 2,958 articles for full-text assessment. After excluding 29 articles for which the full text was unavailable, the remaining 2,929 articles were rigorously evaluated. During this stage, 2,437 articles were excluded as they were not relevant to the core topic or lacked significant, extractable results concerning MSI in cancer. This stringent process resulted in a final cohort of 492 high-quality research articles for data extraction.

**Figure 1.**
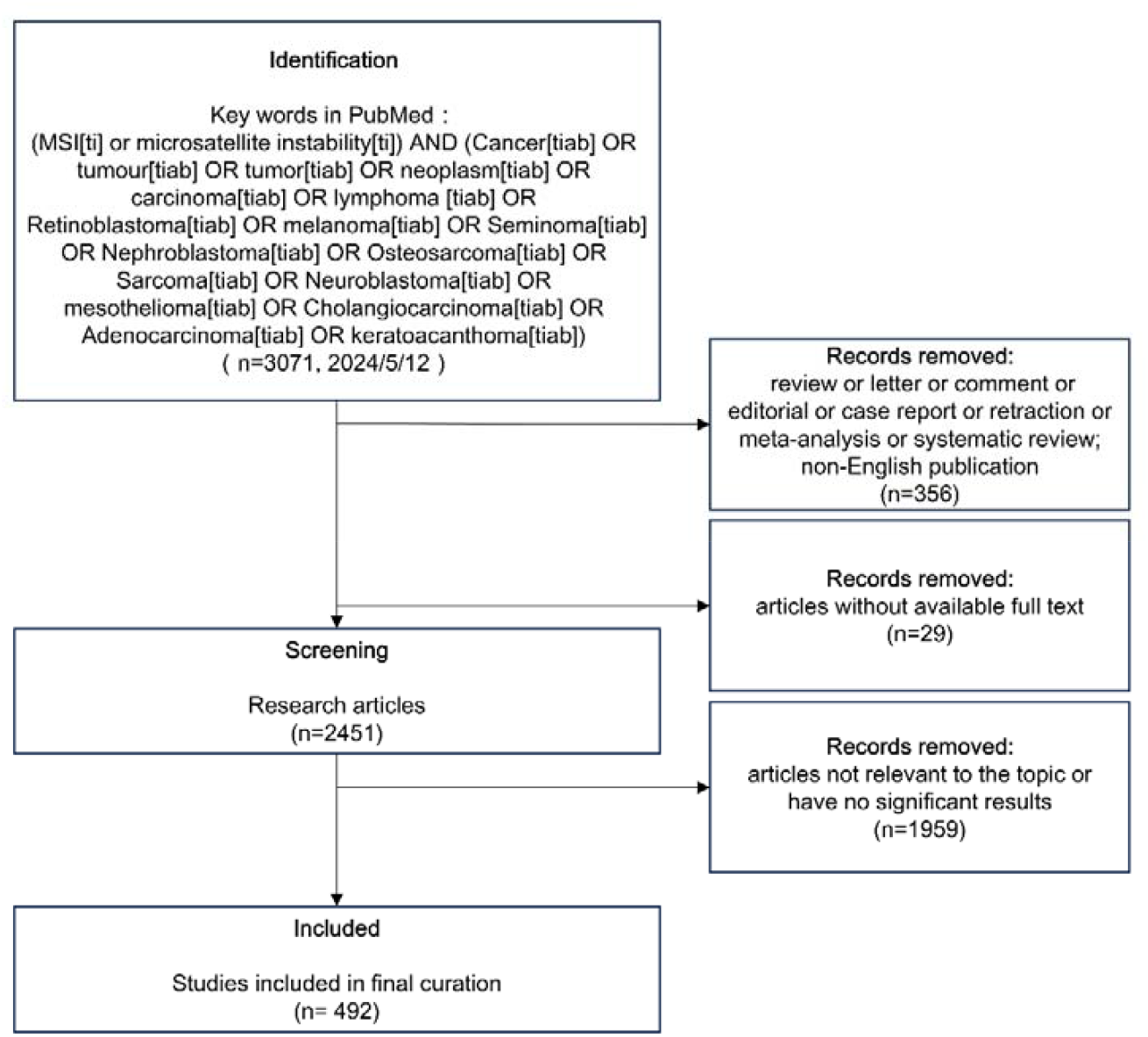
Flowchart of the literature screening and selection process. The diagram illustrates the multi-stage process for identifying relevant studies for inclusion in the MSICKB. Starting from an initial PubMed search, records were systematically screened based on publication type, language, full-text availability, and relevance, resulting in a final cohort of 492 articles for data curation.

### Knowledgebase Architecture and Data Framework

#### Conceptual Design and Schema

Prior to data extraction, we established a robust conceptual framework to systematically organize the multifaceted information related to MSI cancers. Based on a preliminary review of representative literature and considerations of clinical relevance, we defined four primary feature categories: (1) Genetic & Molecular Features (e.g., gene mutations, promoter methylation, gene/protein expression); (2) Clinicopathological Features (e.g., tumor histology, grade, location, patient demographics); (3) Therapeutic Response (e.g., response to immunotherapy, chemotherapy, targeted therapy); and (4) Prognostic Factors(e.g., overall survival, disease-free survival).

The knowledgebase was architected using a relational database model to ensure data integrity, minimize redundancy, and facilitate efficient querying. The core schema comprises six interconnected tables: a Reference table (bibliographic data), a Sample table (cohort details such as cancer type and sample size), and four feature-specific tables corresponding to the aforementioned categories (Figure 2). Foreign keys link the feature tables to the Sample and Reference tables, enabling complex, multi-dimensional queries while maintaining data consistency.

**Figure 2.**
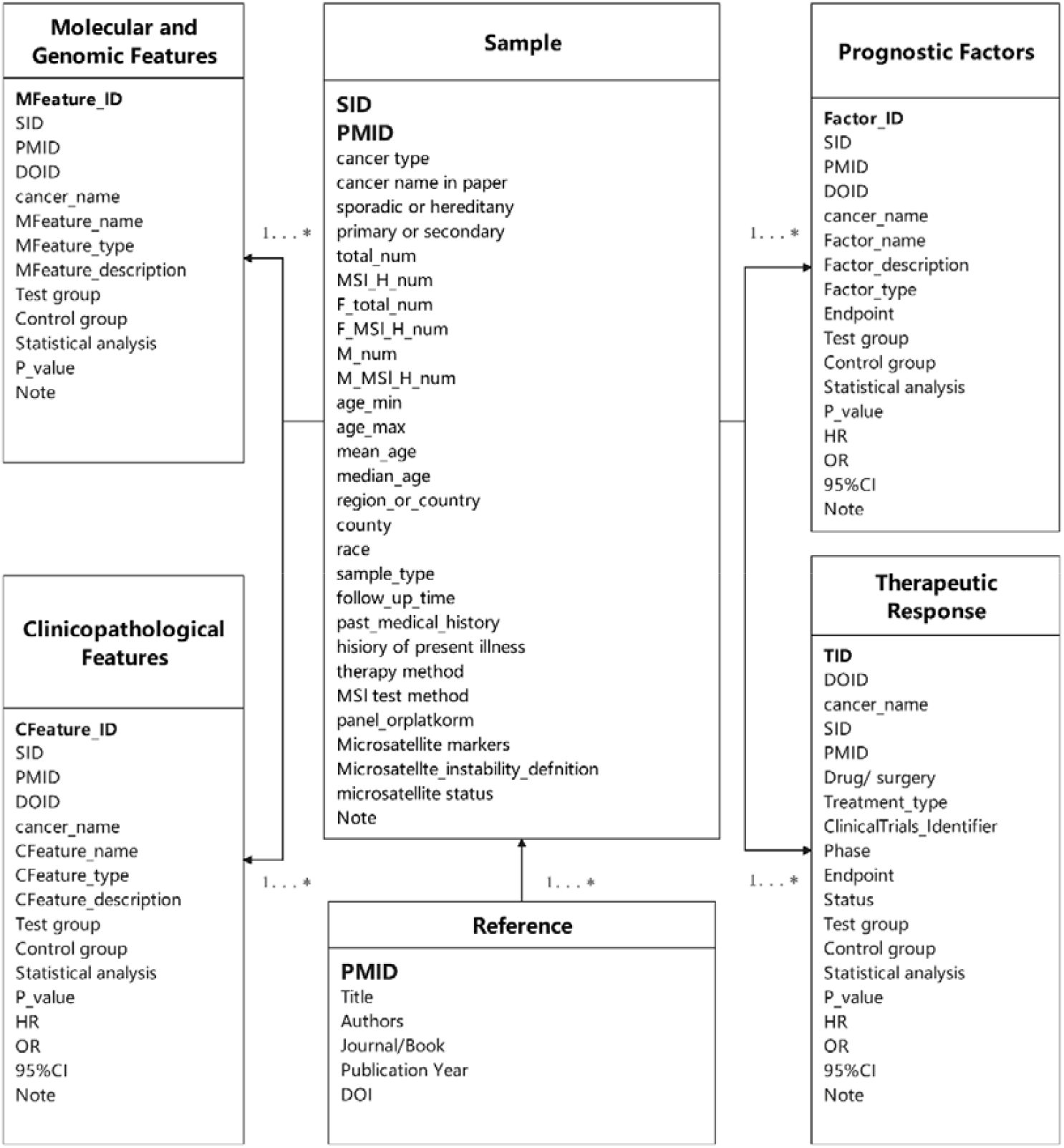
Entity-Relationship (ER) diagram of the MSICKB database schema. The diagram shows the six core tables and their relationships. The Sample and Reference tables serve as central hubs, linked to the four feature-specific tables (Molecular and Genomic Features, Clinicopathological Features, Prognostic Factors, Therapeutic Response). The 1…* notation indicates a one-to-many relationship, where one record in a central table can be associated with multiple records in a feature table.

#### Standardized Data Extraction and Quality Control

To ensure consistency and accuracy, we developed a standardized data extraction protocol based on the predefined schema **(Table 1)**. A rigorous quality control pipeline was implemented. First, all extracted data were standardized using established international guidelines: gene symbols were harmonized according to the HUGO Gene Nomenclature Committee (HGNC) (10), and tumor types were mapped to the Disease Ontology (DO)(11) and the International Classification of Diseases, 11th Revision (ICD-11)(12). Second, a dual-extraction protocol was employed, where two researchers independently extracted data from each article. Any discrepancies were resolved through consensus or consultation with a third senior reviewer. Third, each data entry was assigned a unique identifier for traceability and underwent a final validation check for completeness and accuracy before being integrated into the knowledgebase.

**Table 1:**
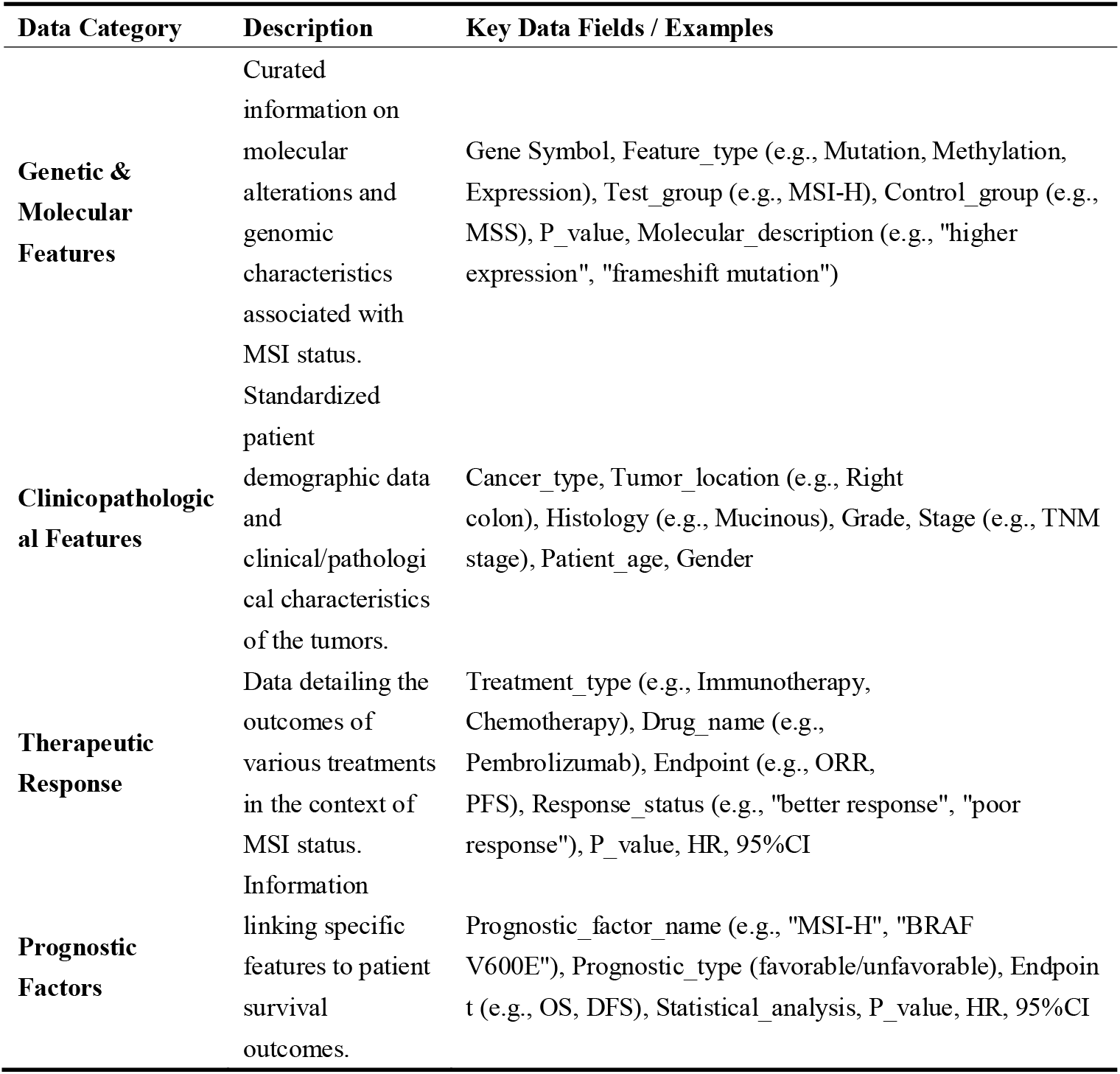
Summary of Data Categories and Key Fields in MSICKB.

### Knowledgebase Implementation and System Architecture

The MSICKB was developed as a web-based platform using the XAMPP stack (v.8.2.12), which integrates Apache (v.2.4.58) as the web server, MySQL (v.8.2.12) as the relational database management system, and PHP (v.8.2.12) for server-side scripting. The front-end user interface was constructed with HTML5, CSS3, and JavaScript, incorporating the Bootstrap framework (v.4.6.2) to ensure a responsive and intuitive user experience across devices. For dynamic data visualization, we integrated the Pyecharts library(v.2.0.7), enabling users to interactively explore the complex relationships within the curated data. To guarantee high performance, especially for complex queries involving multi-table joins, strategic indexing was applied to frequently accessed fields in the MySQL database. This technical architecture ensures that MSICKB is not only a rich data repository but also a powerful and user-friendly platform for scientific inquiry.

### Bioinformatics and Network Analysis

To demonstrate the application of MSICKB for biomarker discovery, we performed a case study on MSI-associated genes and diseases. A gene-disease interaction network was constructed by extracting all gene-disease association pairs from the curated data. The topological properties of this network were analyzed using the Python libraries NetworkX (v3.2.1)(13) and powerlaw (v1.5)(14). The degree distribution was fitted to a power-law model (discrete=True) to assess its scale-free characteristics. The degree threshold for identifying hub genes was determined by examining the degree distribution plot to identify an “elbow point” that separates highly connected nodes from the majority of nodes. Based on this analysis, a degree cutoff of k ≥ 4 was selected.

Hub genes were identified based on a degree cutoff of k ≥ 4. This set of 14 hub genes was then subjected to functional enrichment analysis using Metascape(15), a web-based portal. We analyzed for Gene Ontology (GO) Biological Processes(16,17) and KEGG Pathways(18) to elucidate the collective biological functions of these central nodes. The network was visualized using Cytoscape (v3.10.1)(19), employing a force-directed layout to highlight its structural organization.

## Results

### The MSICKB User Interface and Functionality

#### Website Functionality and User Interface

The MSICKB knowledgebase is accessible via a web portal designed with a dual-access architecture to support both exploratory browsing and hypothesis-driven queries (a schematic of the user interface is provided in **Figure 3A**). For broad, initial exploration, the homepage features browsing modules that allow users to navigate the curated data through pre-defined categories, such as major cancer systems or high-level feature classes (e.g., Genetic, Clinicopathological). This guided-tour approach facilitates user familiarization with the database’s scope and content landscape.

**Figure 3.**
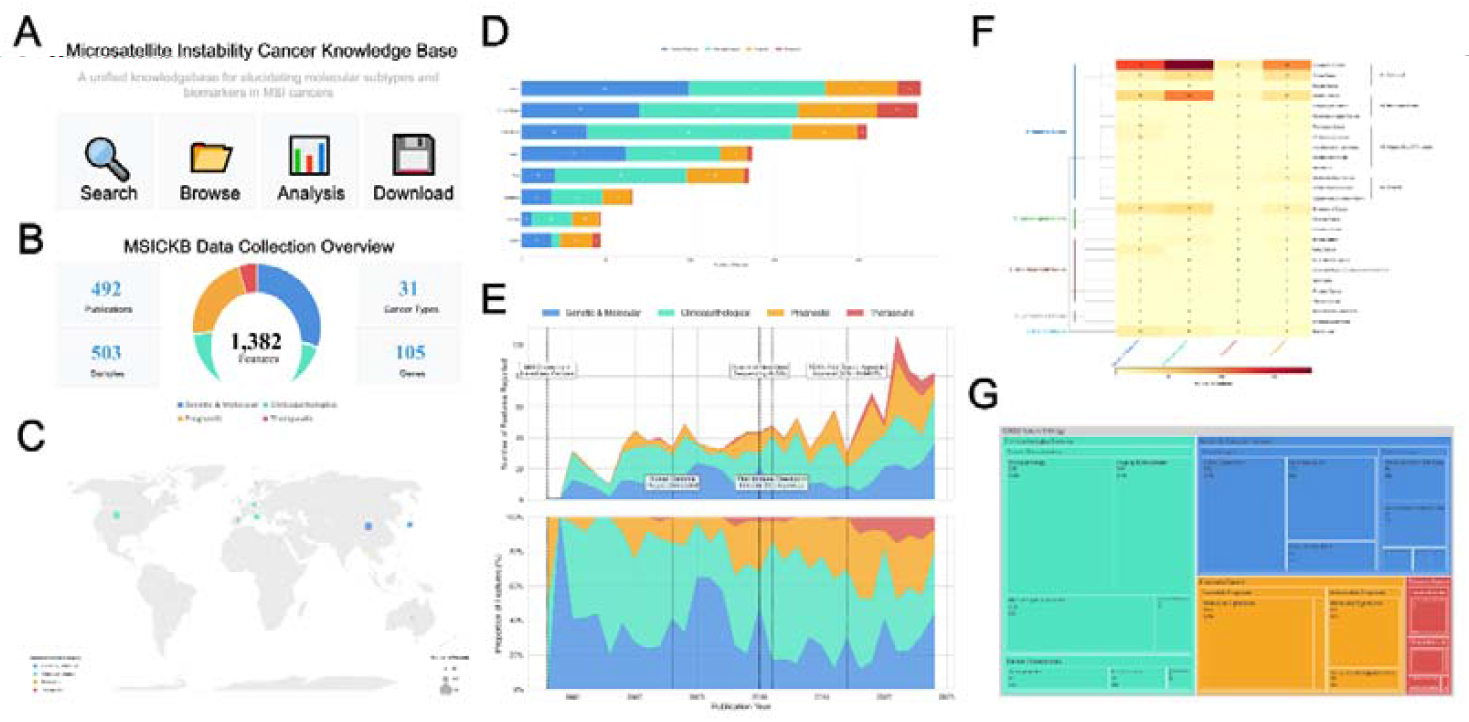
Overview of the MSICKB knowledgebase and data statistics. (A)Schematic of the MSICKB web interface, showing the four main functionalities: Search, Browse, Analysis, and Download. (B)Data collection overview, summarizing the total number of curated publications, samples, cancer types, genes, and the distribution of the 1,382 features across four main categories. (C)Geographical distribution of the included studies. The size of the circle is proportional to the number of features reported from each country, and the color indicates the dominant feature category. (D)Distribution of curated features across the top 8 contributing countries. (E)Temporal trend of MSI-related research from 1995 to 2024. The upper panel shows the absolute number of new features reported each year, annotated with key milestones. The lower panel shows the proportional distribution of the four feature categories over time. (F)Heatmap showing the distribution of the four feature categories across different cancer types, which are hierarchically clustered by organ system. (G)Treemap visualizing the hierarchical ontology of all annotated features in MSICKB, showing the proportion of different subcategories.

For targeted inquiries, a dedicated search engine enables users to perform precise queries using keywords such as a gene symbol (e.g., *TP53*), a cancer type, or a specific clinical term. An advanced search function is also implemented for constructing complex, multi-parameter queries. Search outputs are rendered in a structured, interactive table format, allowing for sorting, filtering, and direct data export. This architectural separation of browsing and searching functionalities ensures that MSICKB serves a wide range of user needs, from exploratory data mining to the retrieval of specific data points, with efficiency and precision.

### Data Content and Statistical Landscape

#### Overall Data Scale and Coverage

The scale and composition of the MSICKB knowledgebase are summarized in **Figure 3B**. The current release synthesizes evidence meticulously curated from 492 peer-reviewed publications, establishing a comprehensive repository of MSI-related cancer knowledge. This curation effort spans 31 distinct cancer types, encompassing data derived from 503 unique samples and documenting molecular and clinical information for 105 individual genes.

Collectively, this multi-source integration has yielded a total of **1**,**382** standardized features, which constitute the core of the knowledgebase. These features are systematically organized into four primary functional categories: **Clinicopathological Features**, which represent the largest portion (44%, n=606); **Genetic & Molecular Features** (26%, n=412); **Prognostic Factors** (20%, n=301); and **Therapeutic Response** data (5%, n=63). This multi-dimensional data architecture, balancing extensive genomic data with clinical outcomes, establishes MSICKB as a robust resource for the systematic investigation of MSI across human cancers.

#### Geographical and Temporal Distribution of Research

To delineate the geographical landscape and historical evolution of MSI-related research, we analyzed the regional origins and publication years of all cataloged features (**Figure 3C-E**).

##### Geographical Distribution

Our data reveal a global research footprint with significant contributions concentrated in specific regions (**Figure 3C**). Features were sourced from studies across 42 countries and territories, with the majority originating from Asia, Europe, and North America. Notably, China and the United States emerge as the two leading contributors in terms of feature volume. An analysis of the top eight contributing countries (**Figure 3D**) highlights their distinct research profiles. For instance, “Genetic & Molecular” studies constitute the largest share of contributions from China (n=99), whereas the United States has made its most significant contribution in the “Clinicopathological” domain (n=94). South Korea exhibits a particularly strong focus on “Clinicopathological” features (n=121), potentially reflecting its strengths in large-scale clinical cohort analyses.

##### Temporal Evolution

The number of reported MSI-related features demonstrates a significant and accelerating upward trend over the past three decades (**Figure 3E**, top panel). This growth has intensified markedly since approximately 2015, a period coinciding with the widespread adoption of next-generation sequencing (NGS) and the landmark FDA approval of immune checkpoint inhibitors (ICIs) for MSI-H/dMMR tumors(8). An examination of the evolving research focus reveals a clear paradigm shift (**Figure 3E**, bottom panel). Early studies were dominated by “Genetic & Molecular” investigations, aligning with milestone events such as the initial discovery of MSI in hereditary cancers(3,20) and the completion of the Human Genome Project(21,22). In recent years, while all categories have grown in absolute numbers, the relative proportion of “Prognostic” and, most notably, “Therapeutic” features have increased substantially. This signals a pivotal shift in the field’s focus from fundamental discovery towards clinical translation and the development of novel treatment strategies.

### Multi-dimensional Data Structure and Hierarchical Ontology

#### Cross-Cancer Feature Landscape

To dissect the multi-dimensional landscape of MSI research across different malignancies, we performed a hierarchical clustering analysis on 31 cancer types based on their feature distribution profiles (**Figure 3F**). This data-driven taxonomy revealed five distinct super-clusters that transcend traditional organ-based classifications. The most prominent cluster is the **Gastrointestinal System (A)**, which is further subdivided into colorectal (A1), esophago-gastric (A2), and hepatobiliary/pancreatic (A3) subgroups, all characterized by a comprehensive and balanced feature profile. Other well-delineated clusters include the **Gynecological Cancers (B)** and **Other Major Solid Tumors (C)**. A salient observation from the heatmap is the pervasive abundance of “Clinicopathological” and “Genetic & Molecular” features across most cancer types, underscoring a primary research focus on the mechanistic and diagnostic aspects of MSI. In contrast, “Therapeutic” and “Prognostic” features are most concentrated in well-studied malignancies like colorectal cancer, thereby highlighting critical knowledge gaps and opportunities for translational research in less-characterized cancers.

#### Hierarchical Annotation Ontology

Complementing the cancer-level analysis, the database’s internal annotation framework organizes all **1**,**382** curated features into a structured, four-level ontology (**Figure 3G**). This hierarchical structure ensures data consistency and facilitates systematic, multi-level querying. At the highest level, the ontology is divided into four principal domains: Clinicopathological Features (n= 606), Genetic & Molecular Features (n= 412), Prognostic Factors (n= 301), and Therapeutic Response (n= 63). Each domain is progressively subdivided into more granular sub-categories. For example, the “Genetic & Molecular Features” domain branches into the “Genetic Layer,” which includes annotations for gene expression, mutation, and methylation, and the “Genomic Layer,” which covers higher-order concepts like MSI status and Tumor Mutational Burden (TMB). This structured ontology is fundamental to the database’s utility, enabling users to navigate from broad biological concepts down to specific, detailed data points with logical precision, thereby enhancing data discoverability and supporting complex bioinformatics analyses.

### Application Example: Analysis of the MSI Cancer Database

#### Construction and Topological Analysis of the Gene-Disease Network

To systematically investigate the relationships between MSI-associated genes and related diseases, we constructed a comprehensive gene-disease network based on the curated data. The resulting network consists of 119 nodes (including 105 genes and 14 diseases) and 297 edges, representing the experimentally validated or reported associations (Figure 4A).

**Figure 4.**
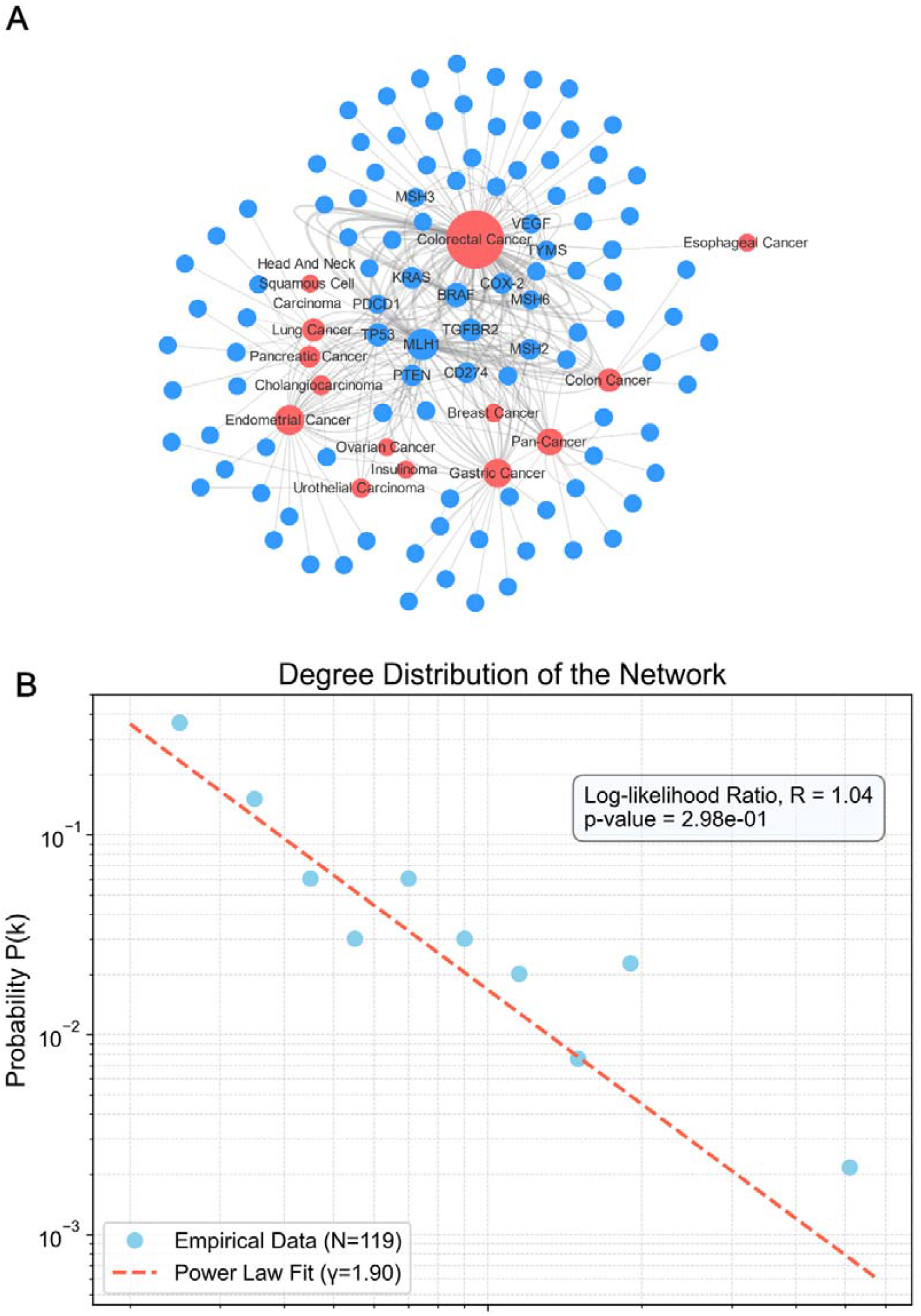
Construction and topological analysis of the MSI gene-disease network. (A) The global view of the gene-disease network, created with Cytoscape. Red circles represent diseases and blue circles represent genes. The size of each node is proportional to its degree. (B) The degree distribution of the network. The plot shows a linear trend on a log-log scale, suggesting scale-free properties. The fitted power-law model (dashed line) yielded an exponent γ = 1.90.

We then analyzed the topological properties of this network. The degree distribution of the nodes was plotted, which displayed a clear linear trend on a log-log scale, a hallmark of networks with scale-free characteristics (Figure 4B). A power-law model was fitted to the distribution, yielding an exponent γ = 1.90. This scale-free-like property suggests that our constructed network is robust and possesses a hierarchical structure. In such a network, the majority of nodes have few connections, while a small number of nodes, known as ‘hub’ nodes, are highly connected. These hubs are often critical for maintaining the network’s structure and function and are presumed to represent key molecular players in the context of MSI cancers.

#### Identification and Functional Enrichment Analysis of Hub Genes

Based on the network’s topological properties, we identified hub genes as those nodes with a degree greater than or equal to 4 (k ≥ 4). This resulted in a list of 14 hub genes (Table 2), which are considered the most central and influential nodes in the MSI-related network.

**Table 2.**
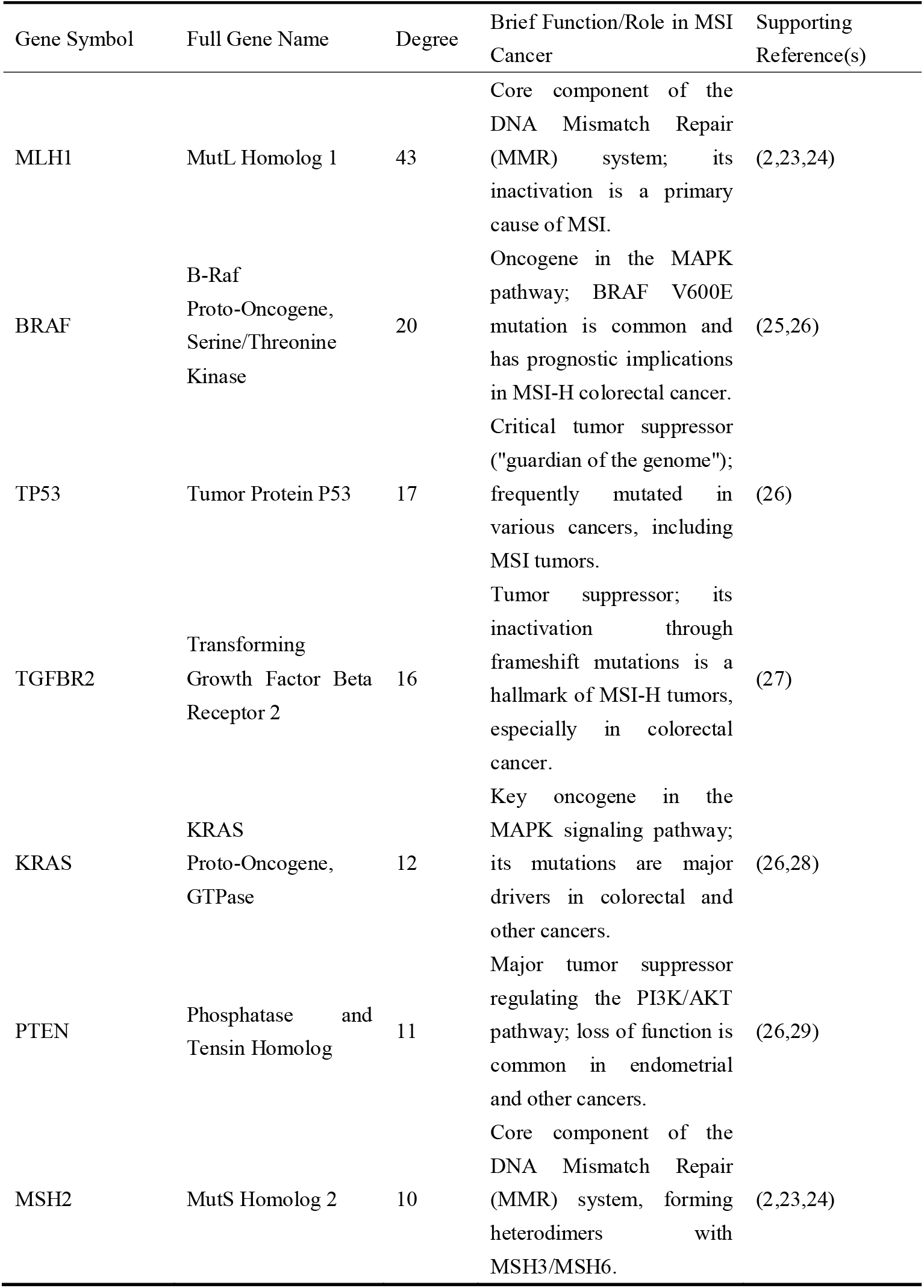
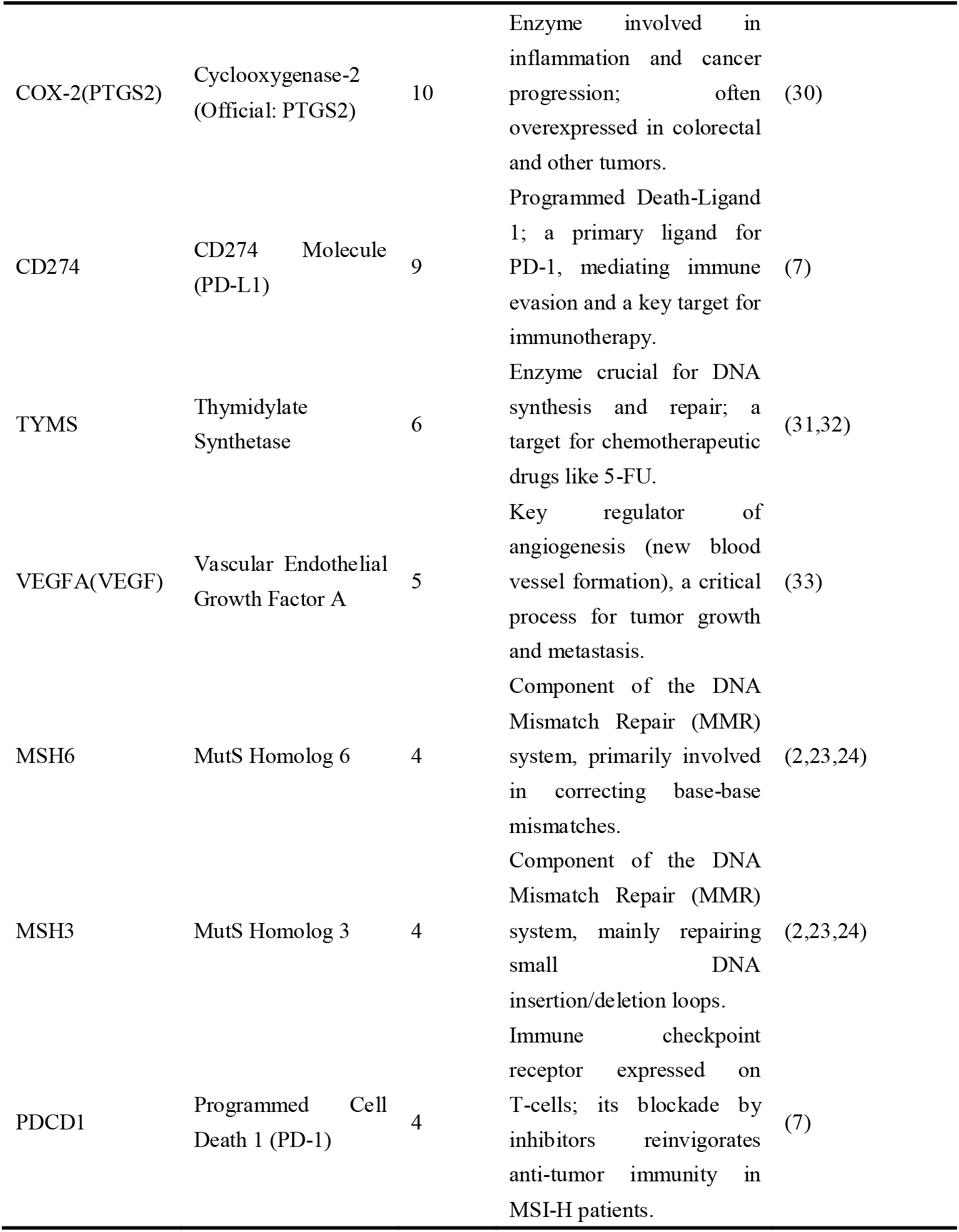
List of the 14 Identified Hub Genes and Their Functions.

To understand the collective biological functions of these 14 hub genes, we performed pathway and process enrichment analysis (Figure 5). The analysis revealed significant enrichment in pathways directly related to MSI, such as **‘Chromosomal and microsatellite instability in colorectal cancer’** (WP4216). Furthermore, the hub genes were significantly enriched in KEGG pathways for key MSI-associated malignancies, including **‘Endometrial cancer’** (hsa05213) and **‘Gastric cancer’** (hsa05226). Crucially, the analysis also highlighted a strong link to immune-related processes, with significant enrichment in terms like **‘regulation of lymphocyte activation’** (GO:0051249), which aligns with the mechanism of action for immune checkpoint inhibitors. These results strongly suggest that the identified hub genes not only are central in the network structure but also functionally converge on the core pathological mechanisms of MSI cancers: DNA repair defects and tumor-immune interactions.

**Figure 5.**
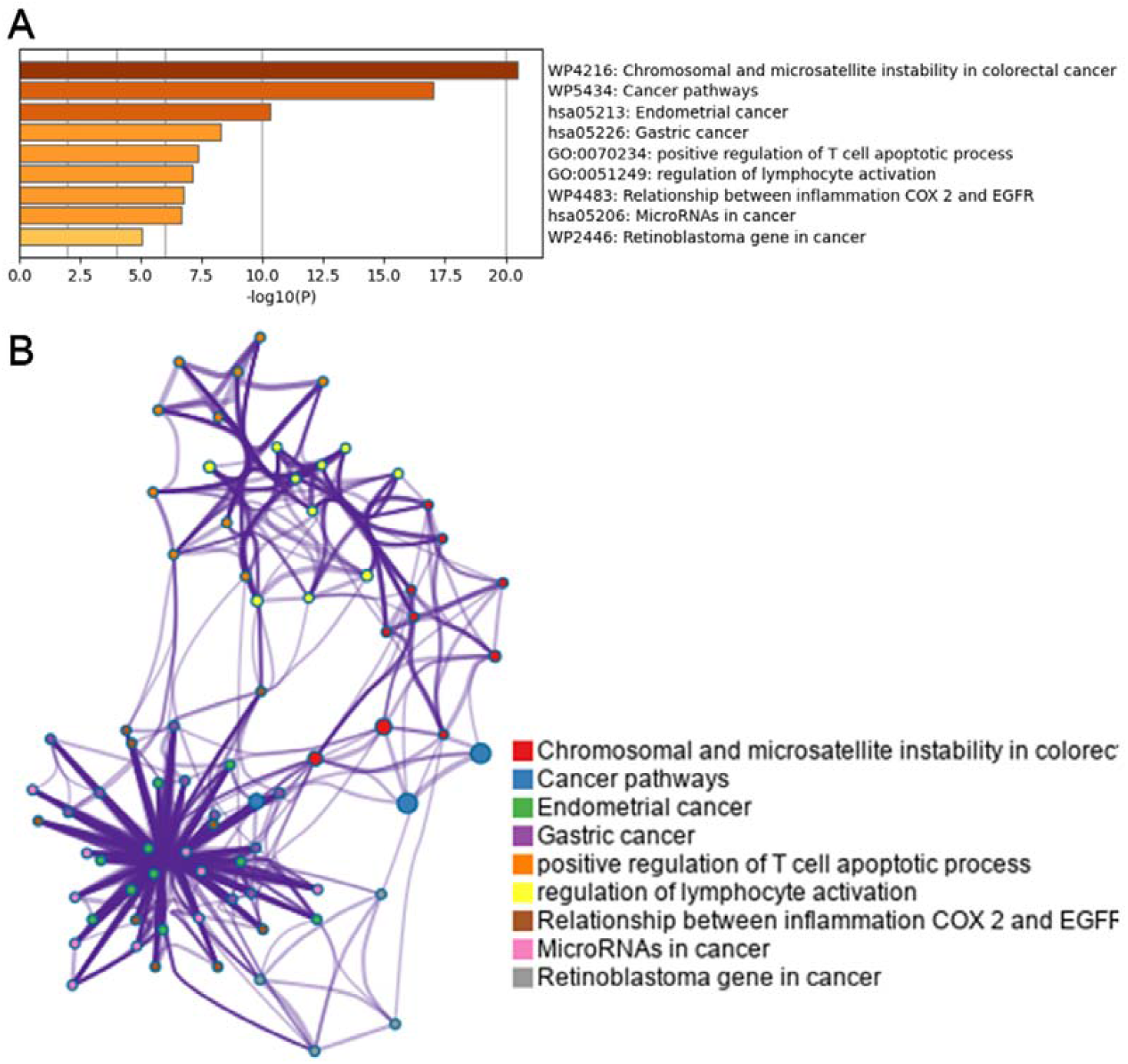
Functional enrichment analysis of the 14 hub genes. (A) Bar chart of the top enriched pathways and processes, ranked by −log10(P-value). (B) Network of enriched terms, where nodes represent enriched terms and are colored by cluster ID. The analysis highlights significant enrichment in pathways related to microsatellite instability, key MSI-associated cancers, and immune regulation.

#### Para.3: Validation of the Reliability of Hub Genes

To further validate the biological significance of the 14 identified hub genes, we examined their established roles in MSI cancers based on existing literature. Remarkably, all 14 genes identified through our unbiased network analysis have been extensively reported to be associated with MSI-related tumorigenesis (as summarized with key citations in Table 2).

For instance, the list is led by the core components of the DNA Mismatch Repair system (**MLH1, MSH2, MSH6, MSH3**), whose deficiency defines the MSI phenotype(2,23,24). It also includes well-established oncogenes and tumor suppressors such as **TP53, PTEN, KRAS, and BRAF**, which are key drivers in cancer progression and frequently altered in MSI tumors(34,35). Most notably, our analysis successfully captured **PDCD1 (PD-1)** and **CD274 (PD-L1)**, the direct molecular targets of immune checkpoint inhibitors, a therapy that has shown profound efficacy in MSI-high patients(7).

The successful identification of this ‘all-star’ panel of genes through a purely topological approach strongly validates the effectiveness of our network-based strategy for pinpointing critical molecular players in complex diseases. The implications of these findings and the broader utility of the MSICKB are discussed below.

## Discussion

In this study, we addressed the critical challenge of molecular heterogeneity in MSI cancers by developing MSICKB, the first comprehensive knowledgebase that systematically integrates multi-dimensional data on microsatellite instability across various cancer types. The application of MSICKB through a network-based analysis provided novel insights into the topological organization of the MSI-associated molecular landscape. Our analysis revealed that the gene-disease network exhibits scale-free-like properties, indicating a hierarchical structure governed by a few highly connected hub nodes(36).

The identification and functional characterization of 14 hub genes represent a key finding of our work. These genes, including core MMR components (e.g., MLH1, MSH2) and pivotal immune regulators (e.g., PD-1, PD-L1), were found to be significantly enriched in pathways central to MSI tumorigenesis, namely DNA repair defects and tumor-immune interactions. The successful recapitulation of this “all-star” panel of known MSI-related genes through an unbiased computational approach not only validates the utility of our knowledgebase but also underscores the power of network analysis in identifying critical biomarkers from complex, curated data(37). These findings provide a solid foundation for future investigations into more refined molecular subtypes and personalized therapeutic strategies for MSI cancers.

While large-scale consortium databases such as TCGA(38) and ICGC(39) have provided an unprecedented foundation for cancer genomics, our MSICKB serves a distinct and complementary role **(Table 3)**. Unlike these repositories, which primarily house raw sequencing and multi-omics data, MSICKB is a knowledge-centric resource built upon manually curated, evidence-based features extracted directly from peer-reviewed literature. This curation process translates complex biological findings into standardized, queryable data points, bridging the gap between raw data and actionable biological knowledge(40–44).

**Table 3.** Comparison of MSICKB with Existing Major Cancer Databases.

| Feature | MSICKB (This work) | TCGA/GDC Portal | cBioPortal | COSMIC |
| --- | --- | --- | --- | --- |
| Primary Data Source | Manually curated from peer-reviewed literature | Raw multi-omics data (e.g., WES, RNA-seq) | Raw multi-omics data (TCGA, ICGC, etc.) | Literature curation & sequencing data |
| Focus | Disease-specific: Microsatellite Instability (MSI) cancers | Pan-cancer genomic and transcriptomic profiles | Pan-cancer genomic data visualization | Pan-cancer somatic mutations |
| Data | Evidence-based, | Gene/protein | Gene-level | Mutation-level |
| **Granularity** | **feature-level knowledge** (e.g., "Gene X predicts poor prognosis in MSI-H CRC") | expression levels, mutation calls, copy number variations | alterations and clinical data | annotations |
| **Core Content Dimensions** | **Integrates 4 key dimensions:**<br>1. Genetic & Molecular<br>2. Clinicopathological<br>3. Therapeutic Response<br>4. Prognostic Factors | Primarily genomic, transcriptomic, and basic clinical data | Primarily genomic and basic clinical data | Primarily somatic mutations and basic clinical data |
| **Therapeutic Data** | **Rich, curated details on drug response**, especially ICIs | Limited and not standardized across studies | Limited; mainly focused on sample metadata | Limited drug resistance mutation data |
| **Prognostic Data** | **Explicitly curated prognostic associations** (e.g., HR, OS, DFS) | Requires re-analysis of survival data | Basic survival plots available | Limited prognostic information |
| **User Accessibility** | **Intuitive web interface with direct search/browse functions** | Data portal for file download and exploration | User-friendly visualization and analysis tools | Searchable web interface |
| **Integrated Analysis Tools** | **Built-in network analysis application example** | Basic visualization tools (e.g., OncoGrid) | OncoPrint, survival analysis, co-expression | None |

The primary advantage of MSICKB lies in its multi-dimensional data integration. It is, to our knowledge, the first database to systematically consolidate genetic, clinicopathological, therapeutic, and prognostic information specifically for MSI cancers into a single, unified framework. This allows researchers to seamlessly investigate cross-dimensional relationships—for example, correlating a specific gene mutation with chemotherapeutic response and overall survival—that would be challenging to assemble using existing, more generalized cancer databases like **cBioPortal**(45) or **COSMIC**(46). The user-friendly web interface and integrated analysis tools further lower the barrier to entry for researchers to explore this complex data landscape.

Despite its comprehensive scope, we acknowledge several limitations of the current version of MSICKB. First, as a literature-based knowledgebase, its content is inherently biased by existing research trends, with well-studied cancers like colorectal cancer having more extensive data than rarer malignancies. Second, the rapid pace of research necessitates continuous updates to maintain the database’s relevance.

Looking forward, these limitations also map out a clear path for future development. We plan to implement a semi-automated curation pipeline to accelerate the integration of new literature and establish a regular update cycle. A key future direction is to integrate MSICKB with large-scale multi-omics datasets to validate our literature-derived findings and uncover novel associations. Ultimately, we envision MSICKB evolving into a dynamic ecosystem that supports the development of clinical prediction models—for instance, leveraging machine learning algorithms to predict MSI status in cancers or ICI response(47,48) based on a combination of genetic and clinical features. By providing this foundational resource, we hope MSICKB will serve as a catalyst for the broader community, accelerating the discovery of novel biomarkers and paving the way for a new era of precision medicine in the management of MSI cancers(49).

## Conclusion

In conclusion, this study addresses the pressing need for a centralized resource to investigate the molecular heterogeneity of MSI cancers by introducing MSICKB, a comprehensive, manually curated knowledgebase. Our work provides not only a rich, multi-dimensional data repository for the scientific community but also demonstrates its utility through a network-based analysis that successfully identified key biological pathways and a panel of validated hub genes. We believe MSICKB will serve as a valuable catalyst for future research, accelerating the discovery of novel biomarkers and ultimately contributing to the advancement of precision medicine for patients with microsatellite instability cancers.

## Data and Code Availability

The MSICKB knowledgebase is publicly accessible at http://www.sysbio.org.cn/MSICKB/. The data analysis scripts used for network construction and topological analysis are available upon reasonable request from the corresponding author.

